# The Constancy of Colored After-Images

**DOI:** 10.1101/095935

**Authors:** Semir Zeki, Samuel Cheadle, Joshua Pepper, Dimitris Mylonas

**Affiliations:** Laboratory of Neurobiology, University College London, Gower Street, London, WC1E 6BT, UK

**Keywords:** color, vision, color constancy, after images

## Abstract

We undertook psychophysical experiments to determine whether the color of the after-image produced by viewing a colored patch which is part of a complex multi-colored scene depends on the wavelength-energy composition of the light reflected from that patch. Our results show that, just as the color of a patch which is part of a complex scene is independent of the wavelength-energy composition of the light coming from it alone, but depends as well on the wavelength-energy composition of the light coming from its surrounds, so is the color of its after-image. Hence, traditional accounts of after-images as being the result of retinal adaptation or the perceptual result of physiological opponency, are inadequate. We propose instead that the color of after-images is generated after colors themselves are generated in the visual brain.

## Introduction

Color constancy refers to our ability to discount the wavelength-energy composition of the light in which a surface or object is viewed and assign a constant color to it. It has been discussed by many authorities on color, including Hermann von Helmholtz (1911), Ewald Hering (1964), William Rushton (1968) and Edwin Land (1974). By color constancy we do not mean that the color of a surface which is part of a complex scene maintains its exact hue, or shade of color, when viewed successively in lights whose wavelength composition is significantly different. The hue will naturally change as the wavelength composition of the light reflected from it and its surrounds changes; it becomes darker or lighter under different illumination conditions, depending upon the predominance of one set of wavebands or another. Hence a better term would be ***constant color category,*** and we use the term constant color to mean constant color categories. But even when the amount of long-wave (red) light reflected from, say, a green surface that is part of a complex scene is twice the amount of middle-wave (green) light, the (green) surface will be perceived as green, and not red (Land, 1974; Zeki, 1983a). In the work reported here, we investigated whether the color of the afterimage produced by viewing a colored surface is also independent of the wavelength-energy composition of the light reflected from it alone but depends as well on the composition of the light coming from the surrounds. If so, this would have a significant bearing on understanding the extent to which the color of the after-image can be accounted for by adaptation or by physiological opponency; it should lead to a new view of how colored afterimages are generated.

There have been several attempts to account for color constancy. Among these have been:

(*a*) Relative adaptation, which von Kries (1905) and others (Helson, Judd & Warren, 1952; Wyszecki & Stiles, 1982; Shapley, 1997) have thought would be retinal and linked to the von Kries formulation of adaptive gains; this theoretically applies some kind of gain to cone cells to maintain the constancy of colors. Others, like Walls (1960), argued that the adaptation is both retinal and non-retinal. Although the adaptation is commonly referred to as “chromatic adaptation”, implying that it is adaptation to a color, it has been usually studied with monochromatic or narrow wavebands of light illuminating a single patch or against a uniform monochromatic surround (Wyszecki & Stiles, 1982; Jameson & Hurvich, 1972), the effects of which might be confined to the retina and its photo-receptors (often referred to as “retinal chromatic adaptation”) or go beyond and occur at some unspecified location in the visual pathways, from retina to cortex. Hence it would more properly qualify as adaptation to specific wavelengths in very reduced conditions, which leaves out of account the dependence of the constancy of colors on the variance of surround colors (Brown & Macleod, 1997);

(*b*) The vaguely defined process of the “unconscious inference” resulting from judgment and learning (Helmholtz, 1911), and (c) memory for colors (Hering, 1964). Both of these imply that higher, cortical, processes are involved.

(*d*) Edwin Land’s (1974) computational theory, based on a hypothetical brain comparison of the lightness record of a scene generated in three different channels, corresponding to the cone channels. This formulation is more neutral as to location of the critical mechanisms, the term “retinex” being based on the supposition that there is a mechanism somewhere between retina and cortex, and possibly distributed between the two, that compares the wavelength composition of light reflected from a surface and from its surrounds in each of the three channels, to generate constant colors.

Viewing of a colored surface has a perceptual consequence, namely the subsequent perceptual appearance of a (negative) colored after image that belongs to a family of colors which is approximately complementary to the one viewed (Burckhardt, 1866; Pridmore, 2008). This perceptual phenomenon was used by Ewald Hering in developing his opponent theory of vision, which Hurvich and Jameson (1957) established on a quantitative basis. Physiological opponency in the visual brain from the retina onwards (Svaetichin, 1956; De Valois, Abramov & Jacobs, 1966; Derrington, Krauskopf & Lennie, 1984; Gouras, 1968; De Monasterio & Gouras, 1975) acts to sharpen the spectral selectivity of chromatic cells, making them more responsive to narrower wavebands of light than the absorption spectra of the three receptors in the retina. Its discovery has played a significant role in accounting for perceptual color opponency, without necessarily discounting an adaptive basis for the observed perceptual phenomenon, along the lines of the von Kries formulations. It also led to the general view that physiological opponency complements receptoral trichromacy, and that color and, by implication, colored after-images are generated by the co-operation of the two in ways yet to be established (Kaiser & Boynton, 1996; Zaidi, Ennis, Cao & Lee, 2012).

The observed physiological wavelength opponency, where cells excited by long-wave light are inhibited by middle-wave light (or vice-versa), and cells excited by short-wave light are inhibited by long-wave plus middle-wave light, is irresistibly close to the documented perceptual color opponency charted quantitatively by Hurvich and Jameson (1957) and others (De Valois et al., 1966; Derrington et al., 1984). Yet the relationship, in terms of action spectra and peak wavelength selectively of cells on the one hand and the observed psychophysical color opponency on the other, is much too loose to enable us to account accurately for the perceptual system in terms of the physiological one, at least up to the primary (area V1) visual cortex (Valberg, 2001; Gegenfurtner & Kiper, 2003).

Because of this loose relationship, we thought it interesting to learn whether perceptual color opponency can be directly related to physiological opponency. We opted to do so within the context of color constancy, by asking whether the color appearance of after images is due to retinal adaptation or whether, like the color of the image itself, it is independent of the precise wavelength-energy composition of the light reflected from it but depends as well on the wavelength energy-composition of the light coming from its surrounds and the ratios between the two (Land, 1986; Land & McCann 1971). If so, it should be, within wide limits, independent of the wavelength-energy composition of the light in which it is viewed (Land, 1974). A convenient approach was to extend Land’s classical Mondrian experiments and ask subjects not only to identify the color of different patches in a Mondrian target, when each was made to reflect roughly the same triplet of energies, but also identify the color of the afterimage produced by viewing each of the patches. This approach, of using a more naturalistic stimulus consisting of many patches of different color illuminated by three broadband sources of light, also had the advantage that it constituted a significant departure from the use of uniform monochromatic patches and surrounds employed by (Anstis, Rogers & Henry, 1978) to investigate whether the color of the after-image is dependent upon “simultaneous color contrast” or “induced colors”. Our question can thus be formally summarised as follows: *Is the color of the after-image of a patch viewed as part of a multicolored scene and illuminated by light of all wavebands due to adaptation to the dominant wavelength reflected from it or is it dependent on its color alone?* As an example, would viewing, say, a green surface that reflects more red light in a more natural context but is perceived as green (color constancy) result in a green after-image, as would be predicted from theories that aim to account for the color of the after-image through retinal chromatic adaptation and physiological wavelength opponency (Rushton & Henry, 1968; Anstis et al., 1978; Craig, 1940; Brindley, 1959; Sakitt, 1976; Virsu, 1978; Hofstoetter, Koch & Kiper, 2004; Williams & Macleod, 1979) or would it result in a red after-image? If the latter, then the implication would be that it is not “chromatic adaptation”, whether retinal or otherwise, and not physiological wavelength opponent mechanisms either, that are the basis of the colored after-image. Rather, it would suggest that the color of the after-image is generated after the colors themselves are generated in the cortex. The work reported here therefore complements earlier physiological work undertaken with single cells in the cortex (Zeki, 1983a).

Throughout, our study is based on the color of the after-image produced by viewing colored patches which are parts of complex multi-colored scenes and which reflect light of many wavebands, rather than on the color of afterimages produced by viewing colored stimuli, either isolated from all surrounds (Williams & Macleod, 1979) or against neutral surrounds (Zaidi et al., 2012), and /or neutral spots against monochromatic surrounds (induction) (Anstis et al., 1978). We made each viewed central patch of the multi-colored display reflect similar triplets of energies belonging to the long, middle and short-wavebands; under these conditions, each patch will maintain its color (color constancy). We asked subjects to report the color of the after-image produced by viewing that patch, in order to determine whether the color of the after-image is, like the color of the image itself, independent of that wavelength composition. If so, then this would lead to the ineluctable conclusion that the colors of afterimages are generated after color themselves are constructed in the visual brain.

## Material and Methods

### Multicolored displays

To allow us to vary the wavelength composition of the light reflected from surfaces without changing their perceived color, we used four Land color Mondrians (see Fig. 1). Each consisted of an arbitrary assembly of rectangular and square patches of different size (Max: 12.5° x 8°; Min: 5° x 4.5°) and color, arranged in such a way that there were no recognizable objects and none of the patches was surrounded by another patch of a single color. The colored patches were made of matte Color Aid papers, which reduced specular reflectance. The color of the surround patches, which extended more than 10° in all directions from the central patch (which subtended 8.25° x 6°), belonged to the family of colors complementary to the central patch (i.e. if the central patch was yellow, the surround patches were in various hues of purple and blue). The spectral power distributions are reported in watts per steradian per square meter (W Sr m^-2^ nm), as in Land’s experiments; they were obtained by measuring the light reflected from each viewed (central) patch with a PR-650 tele-spectroradiometer (see Figure 1). We also show the stimulus specifications for each Mondrian display in 10^°^ relative cone fundamentals (Stockman & Sharpe, 2000), for the central patch and for each of the surrounding patches (see also Table A1 in the Appendix for the numerical values of the cone excitation ratios). We use an extent of 10° from the central patch because psychophysical experiments show that this is the critical spatial range beyond which the modulation of perceived color by its surround rapidly declines (Wachtler, Albright & Sejnowski, 2001). In Figure 1 note that, in the A, B & D displays, the dominant waveband reflected from the central patch was also the dominant waveband reflected from the surround. In display C, the dominant waveband reflected from the surround [green] was the same color as the perceived color [also green] of the afterimage produced by viewing the central [magenta] patch. This arrangement was undertaken to ensure that the color of the after image produced by viewing the patch could not be accounted for by “color induction” produced by the surrounds (Anstis et al., 1978).

**Figure 1.**
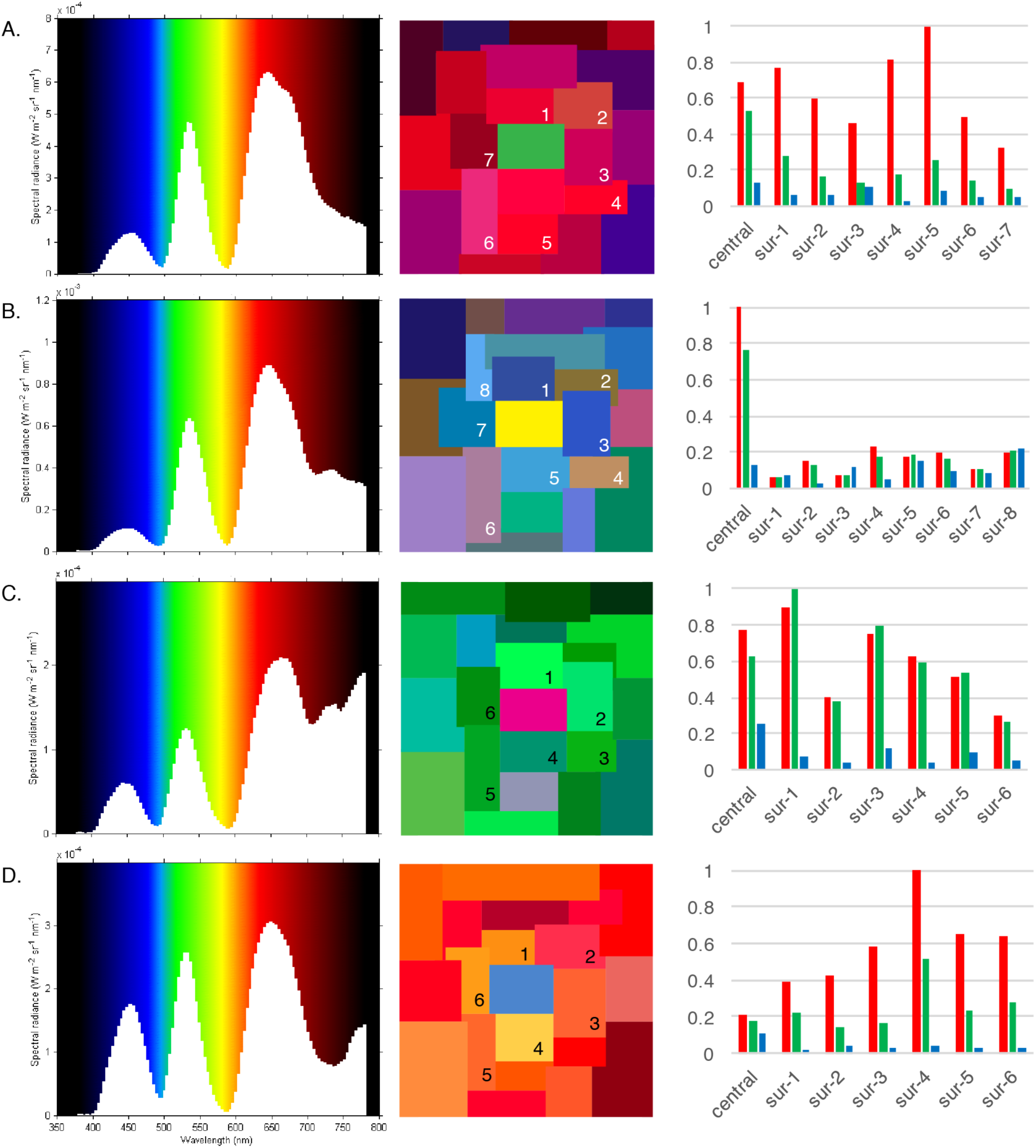
The Mondrian displays used in this experiment and their characteristics. Reflected energies from the four different Mondrian displays (A, B, C & D) (are shown in left the column, color appearance in the center column, and LMS cone excitation ratios in the right column. The spectral power distributions of long (L), middle (M) and short (S) wave light reflected from each central patch (green, yellow, magenta, blue) are given in watts per steradian per metre square per wavelength (W Sr m^-2^ nm). LMS in 10° cone excitation ratios are given separately for the central patch, and for each patch immediately bordering the central patch, up to 10° from the central patch.

### Illumination of the displays

Three 350W Kodak Carousel projectors equipped with rheostats were used to illuminate the Mondrians, as in the original Land Mondrian experiments; each was equipped with specially manufactured long, medium or short wave gelatin filters (Zeki, 1980). Projector 1 transmitted long wave light in the range of 592nm to the end of the visible spectrum with a peak transmittance greater than 660nm; projector 2 transmitted middle wave light in the range 492-580nm (peak 528nm). The short-wave projector transmitted light in the range 386–493nm (peak 432 nm) with a secondary peak at 700nm. Each projector was equipped with a separate rheostat and shutter, thus enabling the intensity of light coming from each to be adjusted separately.

### Color selection targets

To obtain as objective an account as possible of the colors of the after-images, subjects were asked to match the perceived colors of the after images with color patches from the Munsell Book of Color (glossy collection - M40115). Ten approximately uniformly distributed patches of equal value (V=6) and chroma (C=8) were displayed on an annulus at a constant eccentricity of 8° of visual angle. Each patch covered an area of 2° of visual angle. The observers matched the colors of the after images to Munsell samples presented against a white background (see Fig. 4), illuminated with two GrafiLite daylight simulators (CIE 1931 x=0.327, y=0.339, CCT= 5742). The task of the subjects was to determine which patch was the closest match to that of the after image they experienced.

### Observers

Six subjects (5 male; aged between 21 and 35) with normal or corrected to normal vision were recruited using the UCL online subject recruitment system (SONA).

### Adjustments of the visual displays

We adjusted the amount of long, middle, and short wave light reflected from the central (target) patch so that under full illumination conditions (central patch + surrounding patches) it appeared its ‘normal’ color (for example green), even when it was reflecting more light of wavebands that are of the complementary (opponent) family, for example red family in this instance (see Fig. 1). In general, our aim was to make all four central patches reflect the same wavelength-energy composition of light from them while retaining their colors.

### Testing

All four multi-colored displays were placed 2.4m from the projectors and observers sat at a distance of 1.3m from it. The projectors were adjusted appropriately for the particular Mondrian display and set to illuminate the entirety of the display (i.e. both the central patch and the surround). Any remaining light sources in the experimental room were eliminated.

Subjects fixated the central patch of each display for a period of 30s and then reported verbally the color of the after image by choosing one of the four predefined opponent color categories - blue, yellow, red or green. They next selected, from the ten Munsell patches, the one that was the closest match to the color of the after image which they perceived. The color selection targets were placed 60cm away from the observer on a rear desk. The task-lamps were turned on/off immediately after each response to allow the observers to adapt to the viewing conditions of each task while the experimenters were documenting either the verbal or the pointing response. Observers named the color of the afterimages while looking at a white board illuminated identically as in the stimulus presentation phase, and selected the closest Munsell match against a white background. The procedure was repeated three times to measure the reliability of afterimage percepts, giving a total of 72 trials.

We emphasize that, in the natural viewing mode, when a patch was made to reflect more light of a given wavelength, the surrounding patches were so chosen that they, too, would reflect more light of the same wavelength. For example, if a green patch was made to reflect more long-wave light, the surrounds also reflected more long-wave light, even though each patch maintained its color (see Fig. 1). This was to avoid any “induction” effects, as in the experiments of Anstis et al. (1978).

After completing the main experiment observers were tested on the Farnsworth-Munsell 100 Hue test and all were found to be within the normal range (Verriest, Van Laethem & Uvijls, 1982).

## Results

Verbal reports of the colors of afterimages produced by viewing patches that were part of complex abstract colored scenes were grouped into three categories: (I) Reports of opponent colored afterimages are those belonging to the family of colors that is complementary to the color of the target patch (Pridmore, 2008); (II) Reports of non-opponent colored afterimages are those that do not belong to the family of colors that is complementary to the test patch. Examples are those of an afterimage whose color was identical to the test patch, or colored afterimages that bore no clear opponent relationship to the test patch. (III) No response reports were those in which subjects did not report seeing a colored after-image.

The percentage of reports falling into each of the three categories, and averaged across 6 observers, was as follows (see Fig. 2): the most frequently reported color for the after-image belonged to the opponent family (86% of trials), while non-opponent colors were reported in 12.5% of trials. Observers reported perceiving no afterimage in 1.5% of trials. As the primary aim of this experiment was to tabulate the color of the perceived after image, we chose to exclude all no response trials from further analysis. A paired-samples t-test revealed clear evidence that opponent color names were given more frequently than non-opponent color names; t(23)=5.88, p<.0001 for all test displays.

**Figure 2.**
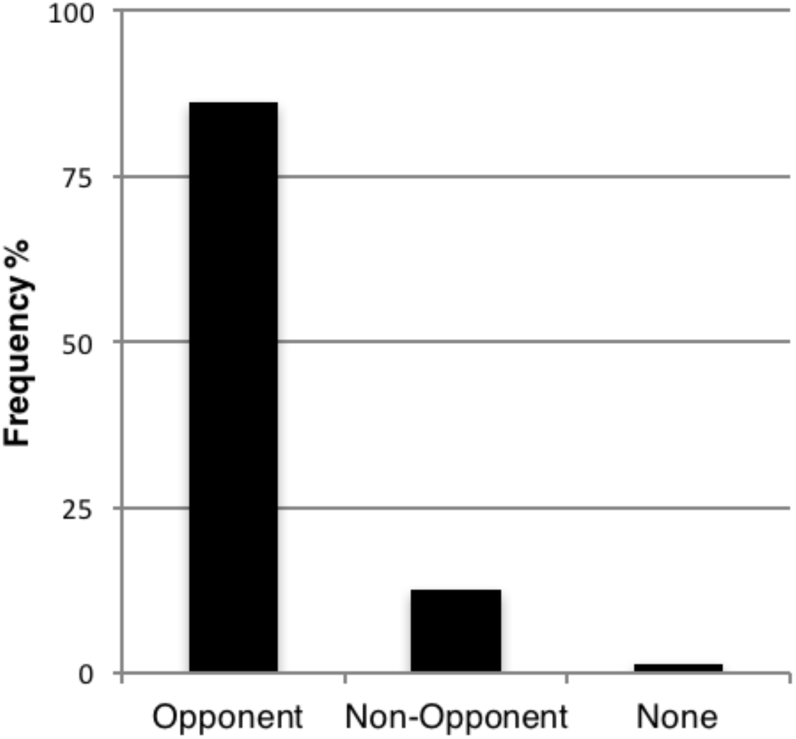
Percentage of responses falling into the three main response categories, averaged across all subjects (n=6).

Figure 3 gives a breakdown of afterimage results, for each of the four display types. Display A (green central patch) produced an opponent after-image in 94% of presentations, display B (yellow central patch) in 100%, display C (magenta central patch) in 100% and display D (blue central patch) in 56%. A paired-samples t-test revealed significant effects for display A; t(5) = 8, p<.0001 and display B; t(5)=17, p<.0001. For display C, we found no variance in the opponent and non-opponent responses – all observers responded an opponent after-image (i.e. 100%). For display D we found no evidence that opponent after-images were given more frequently than not non-opponents (see Discussion); t(5)=.27,p>.05.

**Fig. 3.**
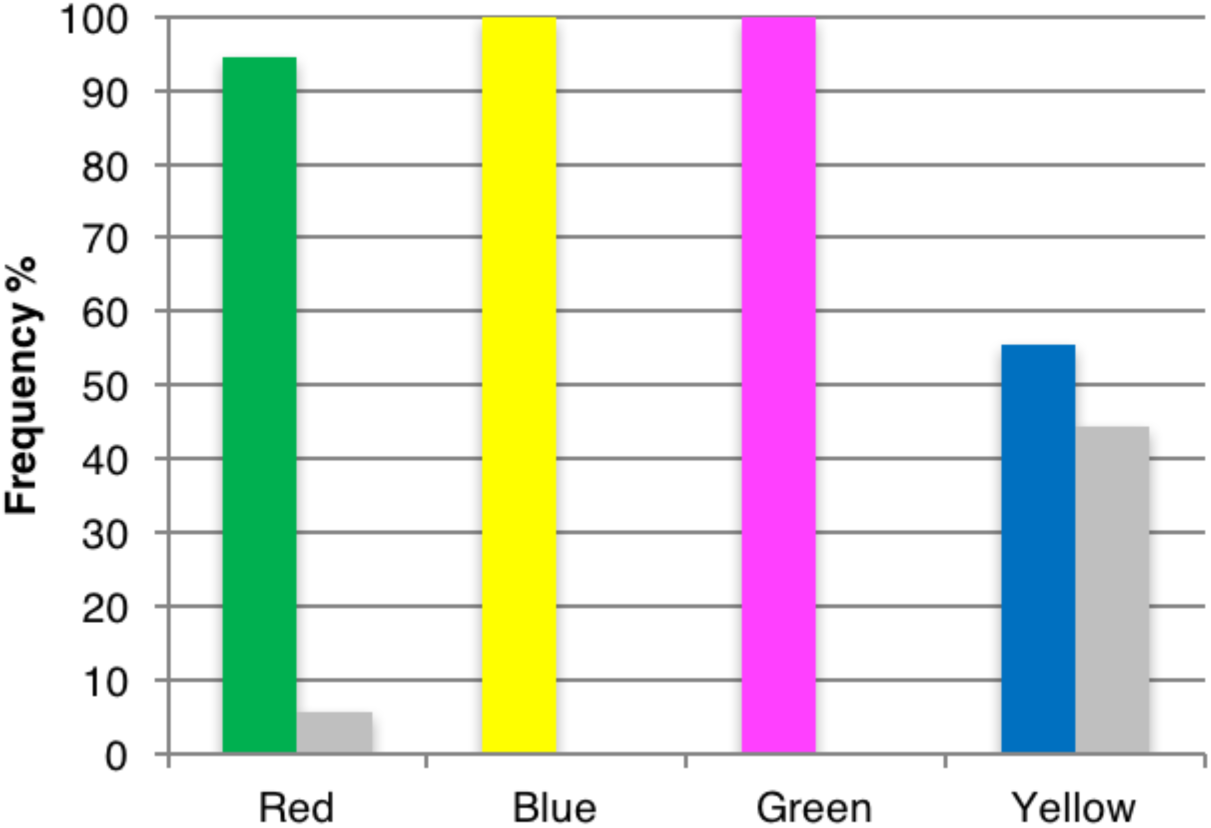
Percentage of opponent and non-opponent colored after-images, for each of the four different display boards used. The color of the bars correspond to the color of the central patches of displays A, B, C and D. Color names indicate the reported color of their After Images. Grey columns correspond to non-opponent responses.

The results demonstrate that, for all display types, opponent colors are perceived more often than ‘all non-opponent’ colors but for display D the effect was not significant.

### Matching the color of the colored after images

The results of the color-matching task were analysed in terms of frequency distributions (see Fig. 4).

**Figure 4.**
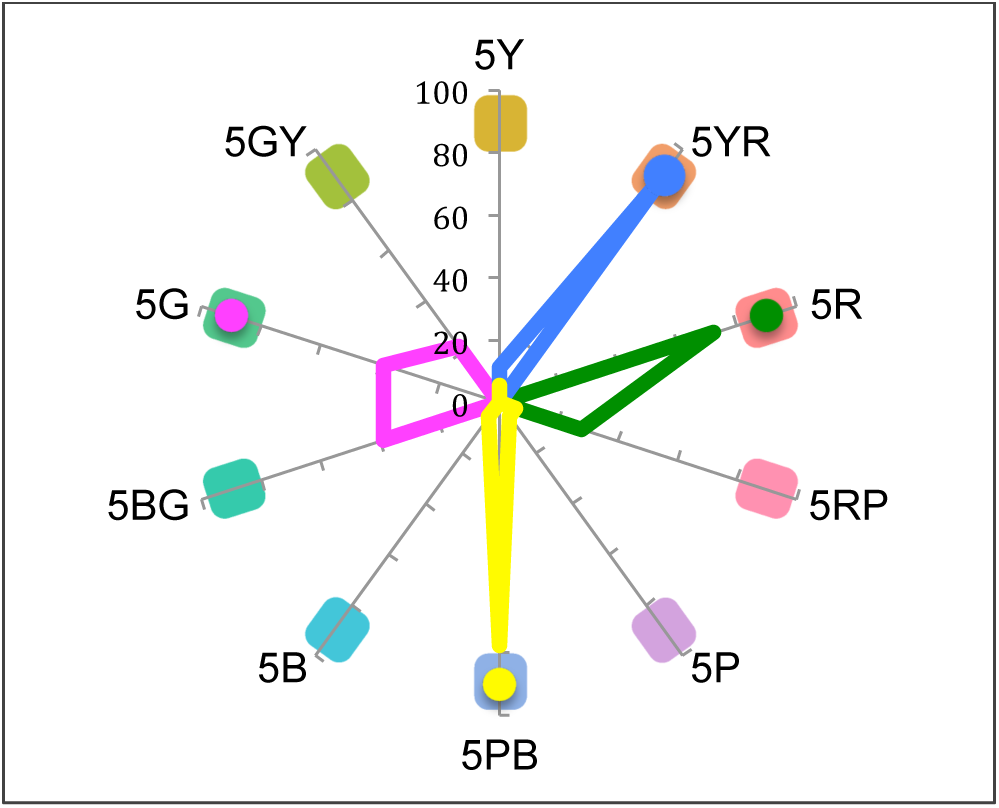
Frequency distributions of color matching tasks are shown for all 4 Mondrian displays. Chip colors and codes (Munsell specifications) are displayed along the polar dimension. Line colors correspond to the color appearance of the central patches (green, yellow, magenta, blue) and dots indicate the median.

The most frequently reported after-images for the green central patch (display A) was red (5R) (72%) while for the yellow central patch (display B) 78% of the observers reported purple-blue (5PB). For the magenta central patch (display C) they reported an afterimage corresponding to blue-green and green (chips 5G and 5BG, 39% for each) and for the blue central patch (display D) they reported yellow-red (5YR) (89%).

In summary, our results thus show that the color of the after-image produced by viewing a colored patch is independent of the precise wavelength-energy composition of the light coming from it, just as the color itself is.

## Discussion

By making each central (viewed) patch reflect the same wavelength-energy composition of light without changing its perceived color (color constancy) we were able to demonstrate that the color of afterimages is opponent to the perceived color, and independent of the wavelength composition of the light (including the dominant wavelength) reflected from the stimulus or from its surrounds alone. The consistency with which the colored after image belonged to the family of colors complementary to the viewed patch was 94% for red, 100% for blue, 100% for green and over 55% for yellow after images. This uniformity of the color naming responses for the after images follows the relative size of these terms (categories) in color spaces (Mylonas & MacDonald, 2016). Since the perceived color is independent of the precise wavelength-energy composition of the light reflected from the viewed patch but depends as well on the wavelength-energy composition of the light reflected from the surrounds (Land, 1974, 1986; Land & McCann 1971), these results demonstrate that the color of the after-image is also independent of the precise wavelength-energy composition of the light reflected from the viewed patch alone.

Although these results are generally consistent with previous reports on the colors of negative after images (Burckhardt, 1866; Pridmore, 2008), there was nevertheless some variability; this was more prominent for the blue central patch which, consistent with previous results (Stromeyer, 1969; Loomis, 1972), produced weaker and more ambiguous after-images. The reasons for this are not clear but they may be related to the relatively lower spatial resolution of the visual system for short-wave (blue) light (Humanski & Wilson, 1992) and their smaller population than L and M cones in the retina, especially in the fovea (Williams, MacLeod & Hayhoe, 1981).

It is difficult to account for these results solely by assuming some kind of retinal von Kries local gain control, because the light of the waveband that was dominant in the light reflected from the central patch was also the dominant waveband in the light coming from the surrounds, up to 10° in all directions, and because afterimages are opponent to perceived colors rather than wavelength, which von Kries mechanisms are assumed to operate on. The lateral extent of the horizontal cells in retina or of cells in V1 do not extend beyond 1-2° for central regions (Ts’o & Gilbert 1988; Packer & Dacey, 2002) and so, to be effective, a von Kries type gain control would have to function over a considerable chain, more extensive than any that has so far been demonstrated. Hence, we agree with previous reports which have questioned the ability of retinal chromatic adaptation and the von Kries rule to fully account for color constancy (West & Brill, 1982; Worthey & Brill, 1986; Foster, 2011; Kulikowski, Daugirdiene, Panorgias, Stanikunas, Vaitkevicius, & Murray, 2012; McCann, & Rizzi, 2012). The relative weak intensity of the light levels of the projectors used in our study (see Fig. 1.) also rules out photochemical effects causing substantial visual pigment bleaching of the photoreceptors (Rushton & Henry, 1968; Williams & Macleod, 1979).

To illustrate the inadequacy of cone contrast mechanisms to explain the formation of after images, we can use as an example Display A (green central patch surrounded by reddish-purple and blue color patches), when the observers projected the colored after-image onto a white board, but the same applies for all viewing conditions. Let us assume that the von Kries rule applies and cones in the retina adapt independently or nearly independently to the central patch of the Mondrian when it is reflecting L, M and S wave light in the ratios of (L=0.684, M=0.530, S=0.134) and to the surround patches (see Fig. 1, 2 and Table A1) in the averaged ratios of (L=0.644, M=0.221, S=0.073). Note that, under these conditions, the L cones are stimulated by more light than M cones. After stimulus offset the mosaic of photoreceptors will generate a set of signals complementary to that generated by the stimulus. Where L, M and S cones had been highly stimulated by the Mondrian they are relatively less responsive to the white board and vice versa. Therefore, the signals sent by the retina would be the same as if looking in a reduction screen setting at the complementary color, greenish and green for proximal field and background area respectively. However, the fact that the color of the afterimage for this display was significantly reported in both tasks as red and not green, as the cone adaptation hypothesis would predict, implies that colored after images are constructed after the signals have left the receptors level in the retina and not until after colors themselves have been generated in the cortex.

The results cannot, as well, be accounted for by physiological wavelength opponency, of the kind demonstrated between retina and cortex (e.g. De Valois et al., 1966). The responses of wavelength opponent cells in V1, for example, have been found to correlate with wavelength composition of the light reaching the eye rather than with its color (Zeki, 1983a). The consequence is that such cells will respond to a surface of any color depending upon the excess, in the light reflected from the surface, of the wavelengths that excite or inhibit it.

The findings we present thus reinforce earlier conclusions which downgrade the importance of cone contrast (Von Kries rule) in the generation of color constancy and assign an increasingly important role to higher cortical mechanisms for the production of colored after images (Murray, Daugirdiene, Stanikunas, Vaitkevicius & Kulikowski, 2006; Rinner & Gegenfurtner, 2002). Collectively, all these results are consistent with a late stage model of afterimage opponency which posits that colored after-images are generated only after the colors themselves are generated in the brain (Zeki, 1983a, 1983b). This is not to say that physiological wavelength opponency mechanisms are not involved in endowing cells with responses that correlate with perceived hues (Zeki, 1983a; Conway, Moeller & Tsao, 2007; Brouwer & Heeger, 2009) but the nature of that involvement remains to be clarified.

The actual cortical site at which after-images are generated is less certain. The results of Delorme (1994) and Tsuchiya & Koch (2005) suggest that positive and negative after images are generated at a binocular level; this makes it plausible to suppose that the level at which colored after images are generated is at a level central to V1. This could be in V2 (Moutoussis & Zeki, 2002) or in V4 (Zeki, 1980). The cells of both are overwhelmingly binocular (Zeki, 1978). V4 is implicated in color vision (Zeki, 1980; Wade, Augath, Logothetis & Wandell, 2008) in both monkey (Zeki, 1983; Wild, Butler, Carden & Kulikowski, 1985; Brewer, Press, Logothetis, & Wandell, 2002; Conway, & Tsao, 2006 and human brains (Wade, Augath, Logothetis & Wandell, 2008 ; Zeki, S., Watson, J. D., Lueck, C. J., Friston, K. J., Kennard, C. and Frackowiak, R. S. 1991; Mckeefry & Zeki, 1997; Goddard, Mannion, Mcdonald, Solomon & Clifford, 2011; Liebe, Logothetis & Rainer, 2011; Lafer-Sousa et al. 2016) and the responses of cells in it can be selective for hues (Zeki, 1980; Conway, Moeller & Tsao, 2007; Brouwer & Heeger, 2013). As well, an imaging study of activity in cortical areas which correlate with the perception of colored after-images pinpointed V4 (Sakai, Watanabe, Onodera, Uchida, Kato, Yamamoto, Koizumi, & Miyashita, 1995). It is of course likely that V4 does not act in isolation but in cooperation with areas V1 and V2, both of which contain sub-compartments of cells which are wavelength selective or opponent and which are reciprocally connected with V4 (Shipp & Zeki, 1995). Although most cells in both V1 and V2 are perhaps more adequately described as wavelength selective, since their responses correlate with the wavelength composition of light as opposed to perceived color Zeki, 1983; Moutoussis & Zeki, 2002), weak surround effects, which may mediate color interactions, have been described in V1 (Wachtler, Sejnowski & Albright, 2003).

### Color and wavelength

Past studies of color vision have been heavily dominated by the use of uniform monochromatically illuminated patches, either in isolation or against monochromatically illuminated backgrounds. Although this has yielded a great deal of cardinal information, it nevertheless has restricted the study of color and of colored after-images to conditions which are not usually encountered in daily life. In more natural viewing conditions, the color of a surface or object is determined by reflection of light of all wavebands from it and from its surrounds, there being a crucial difference in the wavelength-energy composition of the light reflected from the two. It is this difference that ultimately determines color constancy (Land, 1974). Adopting this more natural stimulation condition, we have shown here that, just as perceived colors are constructed by spatial ratio taking operations, so are the perceived colors of the after-image. [We are aware of the tautology implicit in the use of the term perceived color, since there are no colors but perceived colors; we do so only to highlight the point].

We thus make a distinction between adaptation to color when produced by monochromatic light in the usual standard reduction screen setting of a laboratory and when produced by viewing an object or surface that is part of a complex scene when both the surface and its surrounds reflect light of many wavebands. They both produce a colored after-image, but the nature of the colored after-image cannot be adequately studied, or explained, using monochromatic surrounds alone, as done in the studies of Anstis et al., (1978). In particular, it is not possible to tell whether the colored after-image depends on the wavelength-energy composition of the light reflected from a surface or not. It is important to make the distinction between adaptation to color and adaptation to wavelength. We argue that the former is perceptually potent, while the latter, assuming it to exist, is not. This also brings into focus the inadequacy of terms such as “retinal chromatic adaptation”, however much they have come into usage. It perpetuates an historical confusion between wavelength and color (see also Zeki 1983 a and b). While lights of specific wavelength are perceived as having specific colors, the reason for this is traceable to the same laws that operate to generate color, whether attributable to single wavebands of light or to complex configurations where a surface reflects light of all wavebands (Conway et al., 2007). Hence study of after-images using monochromatic light has given an inadequate account of the role of physiological opponency in particular, and color opponency in general, in the generation of color by the cerebral cortex.

In summary, the results of the present study, in conjunction with the evidence described above, indicates that colored afterimages are the result of a late stage mechanism which takes place after colors themselves are generated. The traditionally accepted view of photoreceptor adaptation cannot account for our results and nor can the physiological wavelength opponency observed in the visual pathways from retina to area V2. Our evidence is in favour of a cortical basis for the generation of colored afterimages, one that occurs after colors are generated.

## Acknowledgements

The authors are greatly indebted to John Mollon, Robert Weale, John McCann and Lindsay MacDonald for valuable comments and discussion on earlier versions of this article, and to Peter Zatka-Haas for preparing the Land Mondrian stimuli. This work was supported by the Wellcome Trust, London. Dimitris Mylonas was also supported in part by the UCL Computer Science - EPSRC Doctoral Training Grant, 1573073.

## APPENDIX A

**Table A1.** Summary of long (L), middle (M) and short (S) wave light reflected from the four different Mondrian in 10^o^ cone excitation ratios. LMS ratios are given separately for the central patch, and for each patch immediately bordering the central patch, up to 10^o^ from the central patch.

|  | Target A (green) |  |  | Target B (yellow) |  |  | Target C (magenta) |  |  | Target D (blue) |  |  |
| --- | --- | --- | --- | --- | --- | --- | --- | --- | --- | --- | --- | --- |
|  | L | M | S | L | M | S | L | M | S | L | M | S |
| central | 0.684 | 0.530 | 0.134 | 1.000 | 0.766 | 0.125 | 0.770 | 0.627 | 0.250 | 0.211 | 0.182 | 0.104 |
| surround1 | 0.772 | 0.274 | 0.063 | 0.058 | 0.056 | 0.071 | 0.898 | 1.000 | 0.076 | 0.388 | 0.225 | 0.015 |
| surround2 | 0.599 | 0.160 | 0.063 | 0.154 | 0.123 | 0.026 | 0.399 | 0.381 | 0.035 | 0.430 | 0.141 | 0.041 |
| surround3 | 0.459 | 0.123 | 0.110 | 0.077 | 0.076 | 0.115 | 0.756 | 0.794 | 0.118 | 0.588 | 0.167 | 0.026 |
| surround4 | 0.817 | 0.179 | 0.030 | 0.231 | 0.178 | 0.043 | 0.626 | 0.590 | 0.042 | 1.000 | 0.512 | 0.039 |
| surround5 | 1.000 | 0.255 | 0.087 | 0.178 | 0.187 | 0.153 | 0.512 | 0.533 | 0.095 | 0.651 | 0.231 | 0.033 |
| surround6 | 0.495 | 0.146 | 0.050 | 0.196 | 0.162 | 0.093 | 0.296 | 0.268 | 0.049 | 0.645 | 0.281 | 0.029 |
| surround7 | 0.325 | 0.100 | 0.045 | 0.104 | 0.110 | 0.086 | NA | NA | NA | NA | NA | NA |
| surround8 | NA | NA | NA | 0.201 | 0.211 | 0.215 | NA | NA | NA | NA | NA | NA |

